# Coated crystalline essential amino acid supplementation supports high soybean meal inclusion in diets for Nile tilapia (*Oreochromis niloticus*) with improved protein retention and nutrient assimilation

**DOI:** 10.1101/2024.04.21.590455

**Authors:** Simon J. Davies, Matt E. Bell

## Abstract

The aquaculture industry has previously relied on high-quality fishmeal (FM) to produce diets of excellent standards. However, plant-based proteins such as soybean are more cost-effective for low-value fish species, for example, tilapia and carp, and fishmeal use has been significantly diminished. Previous studies have mainly addressed standard soybean meal (SBM) sources in aquafeeds. A 55-day feeding trial was conducted to investigate the replacement of fish meal with soybean meal concentrate in Nile tilapia (*Oreochromis niloticus)* using a semi-purified diet approach against a Low Temperature (LT) fishmeal as a primary reference protein. Four diets with different levels of soybean meal were evaluated and compared to a control diet of 100 % fish meal. Three diets containing 20, 40, and 80 % soybean protein concentrate (SBPC) were examined. The fourth diet consisted of 80 % SBPC and two essential amino acids: lysine and methionine (80SBPC^AA^) in a coated form. The average daily growth values highlight similar daily growth rates when tilapia was fed 100FM and 40SBPC. No significant differences were observed in the final mean weight for all soybean-fed tilapia, but they were marginally lower than the 100FM control diet group. The 80SBPC diet showed the poorest feed conversion ratio (FCR) at 1.31 and protein efficiency ratio (PER) at 2.19. However, the 80SBPC^AA^ diet showed a significantly improved result for per cent weight gain compared to the 80SBPC-fed tilapia. Also, protein efficiency ratio was significantly higher at 2.47 and had a better Apparent Net Protein Utilization (ANPU) value of 28.51 % compared to un-supplemented 80SBPC-fed tilapia (25.48 %). These results confirmed that high-quality fishmeal can be substituted by up to 40 % SBPC alone and without any detrimental effects on growth or carcass composition. Further studies on plant proteins and supplementary amino acids as a suitable alternative for replacing high-quality fishmeal may promote a more sustainable aquaculture industry.

## 1. Introduction

Contemporary aquaculture encourages researchers to find sustainable alternatives to prolong and improve fish farming. The global demand for fishmeal (FM) is constantly rising and outweighs the industry’s ability to produce it, being a finite resource. With global FM stocks under careful management and scrutiny, cost inflation can threaten the long-term sustainability and profitability of the aquaculture industry. Today’s prime focus is the strategic use of fishmeal in formulated diets for fish, with a focus on alternate sources (Macusi *et al*., 2023). A suitable substitute for FM is paramount, especially in carnivorous fish feeds, but increasing interest also exists for warm water omnivore species that can still include fishmeal. Replacement with suitable plant-based proteins is important, and the aquafeed sector has been using a myriad of such ingredients for many years, but most likely, it is sub-optimally in many scenarios.

Development of feeds for commercially relevant fish species (inc. tilapia) has focussed on protein economy. Additionally, the quality of protein, which contains essential amino acids (EAAs), is crucial for fish growth and development (Kaushik and Hemre, 2008). In this respect, soybean meal (SBM) has been widely applied as a significant substitute for fishmeal in aquafeeds. Obirikorang *et al*. (2020) show the use of SBM reduces the feed cost by 43 % in relation to a commercial FM dependent diet for Nile tilapia (*Oreochromis niloticus*). In addition, SBM has a high-quality protein content, with a steady supply of soybeans worldwide (Sarker *et al*., 2020). Other plant byproducts such as lupins, beans, peas, and pulses show much merit, but soybean continues to be popular on a global scale. The use of commercially processed SBM has proven successful in previous trials for tilapia. Amer *et al*. (2020) stated that FM replacement by 22 % methylated soy protein isolates (MSPI) increases Nile tilapia fingerlings’ final body weight. The authors suggest a maximum replacement of 66 % MSPI without negative effects on tilapia health and performance. Similarly, Zhou and Gu (2020) demonstrated that SBM, when replacing 75 % FM protein, increased Nile tilapia fry final body weight, attaining ∼38.0 g, with a weight gain of over 3,000 %, and impressive Specific Growth Rate, (SGR) of 6.18 % day^-1^. Nevertheless, there is a debate among researchers regarding the benefit of SBM as a protein source for aquafeed for various reasons.

Poor feed efficiency, the presence of antinutritional factors, and negative impact on gut health have been described in some studies (Francis *et al*., 2001; Zhang *et al*., 2018). Additionally, the authors also point out the different methods of processing SBM, diet formulation variation, and differences in fish species and culture systems (Elangovan and Shim, 2000). A higher quality soybean meal arises from more advanced treatment to remove any Anti-Nutritional Factors (ANFs) resulting in a soybean meal concentrate with around 60 % protein content. A common association with SBM is the inadequacy of the essential amino acid profile (particularly, lysine and methionine). However, there is evidence the detrimental effects of soybean replacement are alleviated or perhaps reduced to insignificance with the supplementation of the two essential amino acids believed to be deficient in SBM. For instance, Azarm and Lee (2014) highlight the SBM supplemented with methionine, lysine, and taurine can replace up to 40 % FM in the diet for juvenile black sea bream (*Acanthopagrus schlegeli*). The replacement did not alter fish survival or SGR. Similarly, El-Saidy and Gaber (2002) demonstrated a diet of 55 % SBM supplemented with 0.5 % L-lysine can accurately replace FM in a diet for Nile tilapia fingerlings with no harmful effects on animal performance. These latter authors observed that tilapia fed with the supplemented SBM presented high values of final individual weight and length-specific growth rate (SGR, 1.15), feed conversion ratio (FCR, 1.61), protein efficiency ratio (PER, 0.62) and overall food intake.

More recently, Bell and Davies (2024) reported a study to compare standard-grade soybean meal and corn gluten meal, showing good results with juvenile tilapia assessed by the same metrics. Others have tested soybean meal inclusion with coated amino acids to allow synchronised absorption with intact protein from the intestinal tract to improve their assimilation and post absorption protein biosynthesis. For example, Rajaram *et al*., (2021) tested coated and uncoated crystalline amino acid fortification in formulating a low fishmeal diet for *Penaeus monodon* (Fabricius, 1798). Their findings suggested that coated amino acids proved superior with respect to growth, digestibility, body composition, haemolymph indices and nitrogen metabolism in shrimp.

The present study evaluates incremental inclusion levels of high-quality soybean protein concentrate meal (SBPC) at the expense of fishmeal with the addition of coated essential amino acids (EAAs) lysine and methionine at the highest inclusion level of 80 % fishmeal substitution. The aims were to assess the effects of increasing levels of SBPC on carcass composition and nutrient retention in the Nile tilapia. Additionally, the present research study assesses the effects of supplementing a coated form of lysine and methionine crystalline amino acids on fish performance. Additionally, our study determines whether EAAs-supplemented SBPC could be an economically viable FM replacement alternative in practical feeds for tilapia towards achieving lower fishmeal-formulated diets.

## 2. Materials and methods

### 2.1 Diet design and formulation

A total of five test diets were produced for the feed trial (Table 1). The basal control contained 100 % FM as the main protein source. The further four diets were formulated with levels of 20 % (20SBPC), 40 % (40SBPC), and 80 % (80SBPC) FM being replaced with SBPC. The final diet consisted of 80 % FM and an additional two essential amino acids: methionine and lysine. This is to examine whether crystalline amino acid sources could mitigate the effects of increase SBPC inclusion. All diets were formulated to be isonitrogenous (37 % protein) and isolipidic (15 %). Test diets were produced by a single screw bench extruder using a 2 mm extrusion die and diet mixture preparation as described by Bell and Davies (2024).

**Table 1:** Test diet formulations for Nile tilapia (*Oreochromis niloticus*) and their corresponding proximate compositions (%, DW).

| Ingredient | 100FM | 20SBPC | 40SBPC | 80SBPC | 80SBPC<sup>AA</sup> |
| --- | --- | --- | --- | --- | --- |
| Fish meal<sup>1</sup> | 45.00 | 36.00 | 27.00 | 9.00 | 9.00 |
| Soya protein concentrate<sup>2</sup> | 0.00 | 9.81 | 19.64 | 39.27 | 39.27 |
| Wheat meal<sup>3</sup> | 25.00 | 20.00 | 20.00 | 19.00 | 17.20 |
| Blood meal<sup>4</sup> | 2.00 | 2.00 | 2.00 | 2.00 | 2.00 |
| Dextrin<sup>5</sup> | 5.00 | 10.00 | 10.00 | 10.00 | 10.00 |
| Potato starch | 2.50 | 5.00 | 5.00 | 5.00 | 5.00 |
| Oil (fish: corn oil, 3:1)<sup>6</sup> | 10.05 | 10.94 | 11.83 | 13.33 | 13.33 |
| Vitamin mix<sup>7</sup> | 2.00 | 2.00 | 2.00 | 2.00 | 2.00 |
| Mineral mix<sup>7</sup> | 1.00 | 1.00 | 1.00 | 1.00 | 1.00 |
| Carboxy-methyl cellulose<sup>5</sup> | 2.00 | 2.00 | 2.00 | 2.00 | 2.00 |
| α-cellulose<sup>5</sup> | 6.35 | 4.35 | 2.33 | 0.00 | 0.00 |
| Mono-calcium phosphate<sup>5</sup> | 1.60 | 1.90 | 2.20 | 2.40 | 2.40 |
| L-methionine<sup>5</sup> | - | - | - | - | 0.90 |
| L-lysine<sup>5</sup> | - | - | - | - | 0.90 |
| **Proximate Composition** |  |  |  |  |  |
| Moisture | 11.99 | 11.73 | 11.57 | 11.48 | 11.77 |
| Protein | 33.68 | 34.36 | 34.20 | 34.89 | 33.52 |
| Lipid | 15.65 | 15.69 | 15.40 | 15.53 | 14.96 |
| Ash | 7.36 | 6.88 | 6.66 | 5.50 | 4.94 |

Diet designation code: FM (Fish Meal), SBPC (Soybean Protein Concentrate meal), SBPC<sup>AA</sup> (SBPC supplemented with methionine and lysine).
<sup>1</sup>Fishmeal: LT-94, Skretting, Norway.
<sup>2</sup>Central Soya Ltd, Michigan USA
<sup>3</sup>Kalpro S<sup>TM</sup> Orans, Paris, France.
<sup>4</sup>American Protein Corporation, Des Moines, Iowa, USA.
<sup>5</sup>Sigma-Aldrich, St. Louis, Missouri, USA.
<sup>6</sup>Sevenses, Hull, UK
<sup>7</sup>PNP, UK.

### 2.2 Experimental procedure

400 male tilapia (*O. niloticus*) fingerlings (first generation all XY from YY male, GMT and XX female tilapia) were supplied by FishGen Ltd (Camberley, England) and acclimated for two weeks in circular 40 L tanks that were part of a recirculating aquaculture system (RAS). Before the trial, fish were fed with a commercial diet (Skretting, Norway). The flow rate into each tank was ∼4 L min^−1^, with the overall RAS kept at pH 7-8 and 28 °C throughout the trial. The photoperiod was set at 14:10, light and darkness.

Within this trial, 335 male tilapias were used, with 20 fish randomly assigned to each triplicate tank per treatment. The feeding trial was carried out over 55-days, and tanks were fed 3 % of the total biomass per tank twice a day in two equal portions (1000 and 1700). Fish weights were measured every ten days after feed deprivation for 24 hrs to allow gut clearance, and the feed/body weight ratio was adjusted to reflect the weight gain increase every 10 days. For initial carcass analysis using terminal anaesthesia (MS222) followed by cranial percussion and destruction of the brain, 20 fish were culled. Also, at the conclusion of the trial, five randomly selected fish from each tank were again euthanised (MS222) and cranial percussion with destruction of the brain to confirm death for final carcass analysis.

### 2.3 Proximate analysis

Crude protein, lipid, moisture, and ash of diets and fish body samples were determined using standardised methods by the Official Methods of Analysis (AOAC, 2023). Samples were homogenised using a blender before analysis commenced. Moisture content was determined by heating samples at 105°C for 48h. Protein determination was carried according the Kjeldhal procedure by acid digestion and sequential distillation (Gerhardt Vapodest 3S, Königswinter, Germany). Crude protein content was calculated from the recovered Nitrogen using a 6.25 conversion factor (N*6.25). Crude lipid concentrations were determined using the Folch *et al*., (1957) method using chloroform and methanol extraction (2:1). For ash, the values were calculated by the percentage weight difference after samples being ignited at 550°C for 16h in a muffle furnace until a constant weight is achieved as dried ash. Proximate analysis of diets is presented in Table 1.

### 2.4. Growth performance and feed utilisation indices

The following indices are used as a measure of fish growth performance and feed utilisation during the feed trial:

Mean Weight Gain = final mean weight (g) – initial mean weight (g)

ADG – Average Daily Growth = Total growth (g) / Number of feeding days

SGR – Specific Growth Rate = ((Ln final mean weight (g)− Ln initial mean weight (g)) / days fed) * 100

FCR – Food Conversion Ratio = feed intake (g) / weight gain (g)

ANPU – Apparent Net Protein Utilisation = ((Final retained protein (g) – Initial retained protein (g)) / Protein intake (g)) * 100

PER – Protein Efficiency Ratio = Weight gain (g) / Protein intake (g)

### 2.5. Statistical analysis

Data values are expressed as mean values with their corresponding standard error. Datasets were analysed using one-way analysis of variance and statistical differences were discerned using a post-hoc Tukey test. Statistical significance was considered when P<0.05.

## 3. Results

### 3.1 Growth performance

Table 2 presents the growth performance of tilapia over the 55-day feeding trial. The ADG values showed that 100FM-fed fish had the highest average daily growth rate (0.68 g fish^-1^ day^-1^) but the 40SBPC diet also performed equally well (0.62 g fish^-1^ day^-1^). The 80SBPC diet without EAAs supplementations yielded the lowest ADG value (0.56 g fish^-1^ day^-1^). The results indicated that the relationship between SGR and FCR is inversely proportional. Hence, the lowest value for the FCR was coupled with the highest rate of growth, indicating the most efficient use of food at this level. The 80SBPC-fed tilapia displayed numerically the highest FCR (1.31) for any treatment group. However, supplementation with essential amino acids methionine and lysine resulted in an approximate 8 % significant decrease or better FCR of the 80SBPC^AA^ diet group 1.21). The tilapia receiving the 40SBPC diet has the highest APNU (Apparent Net Protein Utilisation) of 31.25 %. The 80SBPC diet recorded the lowest value for Apparent Net Protein Utilization (ANPU) at 25.48 %. Likewise, PER confirmed 40SBPC to have a higher value of 2.57, with the 80SBPC diet recording the lowest 2.19 and significant. The amino acid supplementation (80SBPC^AA^) gave a 12 % increase in PER in relation to the 80SBPC diet, which correlates to the same increase when measuring the ANPU performance metric.

**Table 2:**
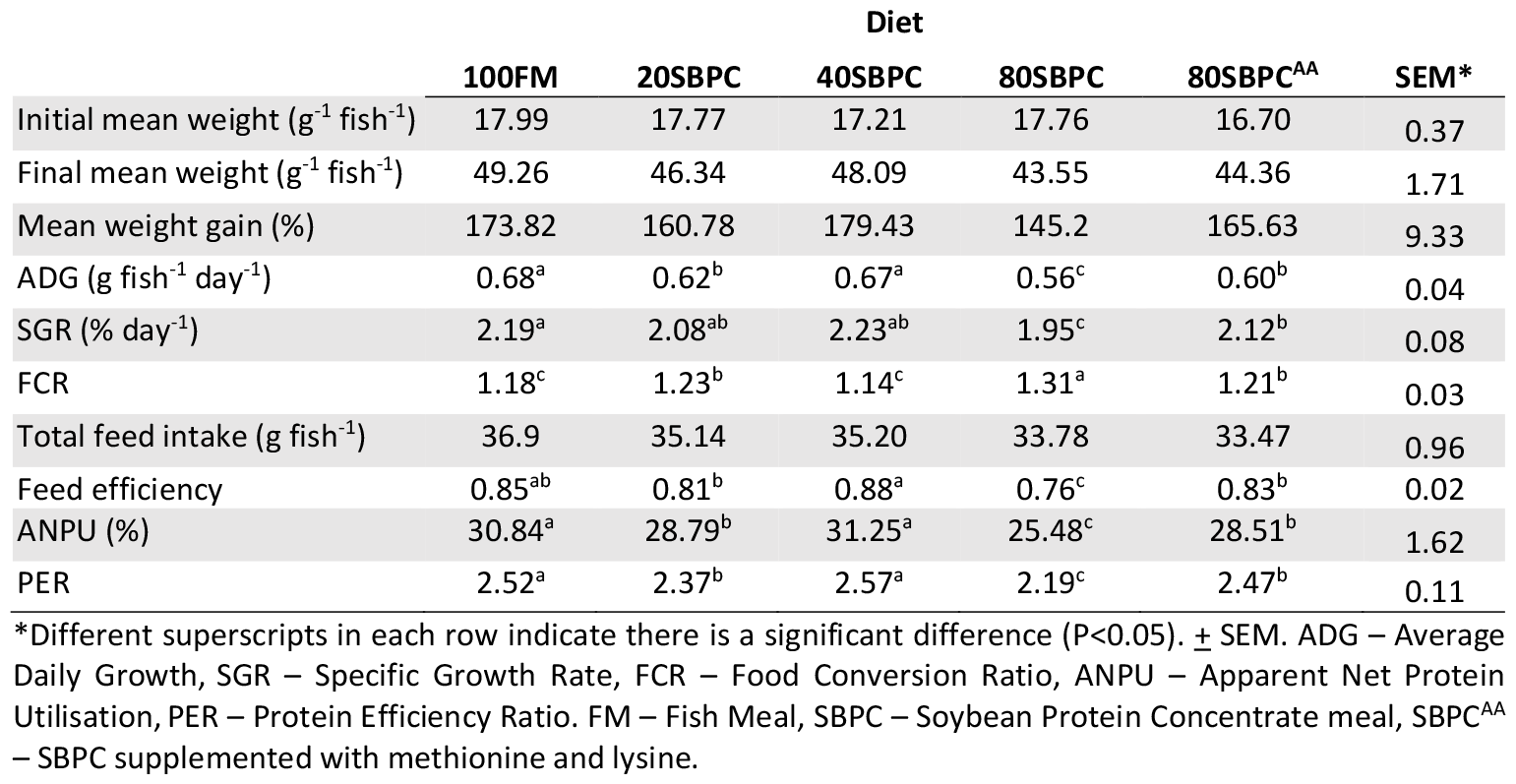
Growth performance and feed utilisation of Nile tilapia (*Oreochromis niloticus*) after being fed increasing soybean protein concentrate meal and animal acid supplementation (Mean values +/-Pooled Standard Error of Mean, SEM).

|  | Diet |  |  |  |  | SEM* |
| --- | --- | --- | --- | --- | --- | --- |
|  | 100FM | 20SBPC | 40SBPC | 80SBPC | 80SBPC<sup>AA</sup> |  |
| Initial mean weight (g<sup>-1</sup> fish<sup>-1</sup>) | 17.99 | 17.77 | 17.21 | 17.76 | 16.70 | 0.37 |
| Final mean weight (g<sup>-1</sup> fish<sup>-1</sup>) | 49.26 | 46.34 | 48.09 | 43.55 | 44.36 | 1.71 |
| Mean weight gain (%) | 173.82 | 160.78 | 179.43 | 145.2 | 165.63 | 9.33 |
| ADG (g fish<sup>-1</sup> day<sup>-1</sup>) | 0.68<sup>a</sup> | 0.62<sup>b</sup> | 0.67<sup>a</sup> | 0.56<sup>c</sup> | 0.60<sup>b</sup> | 0.04 |
| SGR (% day<sup>-1</sup>) | 2.19<sup>a</sup> | 2.08<sup>ab</sup> | 2.23<sup>ab</sup> | 1.95<sup>c</sup> | 2.12<sup>b</sup> | 0.08 |
| FCR | 1.18<sup>c</sup> | 1.23<sup>b</sup> | 1.14<sup>c</sup> | 1.31<sup>a</sup> | 1.21<sup>b</sup> | 0.03 |
| Total feed intake (g fish<sup>-1</sup>) | 36.9 | 35.14 | 35.20 | 33.78 | 33.47 | 0.96 |
| Feed efficiency | 0.85<sup>ab</sup> | 0.81<sup>b</sup> | 0.88<sup>a</sup> | 0.76<sup>c</sup> | 0.83<sup>b</sup> | 0.02 |
| ANPU (%) | 30.84<sup>a</sup> | 28.79<sup>b</sup> | 31.25<sup>a</sup> | 25.48<sup>c</sup> | 28.51<sup>b</sup> | 1.62 |
| PER | 2.52<sup>a</sup> | 2.37<sup>b</sup> | 2.57<sup>a</sup> | 2.19<sup>c</sup> | 2.47<sup>b</sup> | 0.11 |

\*Different superscripts in each row indicate there is a significant difference (P<0.05). ± SEM. ADG – Average Daily Growth, SGR – Specific Growth Rate, FCR – Food Conversion Ratio, ANPU – Apparent Net Protein Utilisation, PER – Protein Efficiency Ratio. FM – Fish Meal, SBPC – Soybean Protein Concentrate meal, SBPC<sup>AA</sup> – SBPC supplemented with methionine and lysine.

### 3.2 Body Composition

The values obtained for moisture ranged from 72.0-72.65 % of the total biomass and were not deemed to be significantly different (P>0.05) (Table 3). The protein content had an inversely proportional relationship to the soybean content of the diet (15.71, 15.63, 15.51, 15.33, 15.38 %), but despite this obvious trend, it was not found to be significant (Table 3). No significant differences in the carcass lipid concentrations were also noted (P>0.05). The ash content of tilapia progressively decreased (4.23, 4.16, 4.12 %) with an increasing soybean meal substitution (0-40 %) but was insignificant (Table 3). However, the 80SBPC and 80SBPC^AA^ diets caused a further significant decrease (3.88-3.84 %) in the whole-body carcass of tilapia compared to the other diet groups (P<0.05) (Table 3).

**Table 3:** Proximate composition of the pooled carcasses of test animals presented as a proportion of the whole fish (%).

|  | Initial | 100FM | 20SBPC | 60SBPC | 80 SBPC | 80SBPC<sup>AA</sup> | *SEM |
| --- | --- | --- | --- | --- | --- | --- | --- |
| Moisture | 64.79 | 72.42 ±0.22 | 72.07 ± 0.19 | 72.01 ± 0.06 | 72.65 ± 0.08 | 72.35 ± 0.10 | 0.19 |
| Protein | 19.68 | 15.71 ± 0.13 | 15.63 ± 0.08 | 15.51 ±0.12 | 15.33 ±0.14 | 15.38 ±0.13 | 0.11 |
| Lipid | 10.53 | 8.35 ± 0.16 | 8.45 ± 0.18 | 8.78 ± 0.08 | 7.99 ± 0.29 | 8.77 ± 0.55 | 0.23 |
| Ash | 5.77 | 4.23<sup>a</sup> ± 0.1 | 4.16<sup>a</sup> ± 0.04 | 4.12<sup>a</sup> ± 0.06 | 3.88<sup>b</sup> ± 0.11 | 3.84<sup>b</sup> ± 0.05 | 0.12 |

\*Different superscripts in each row indicate there is a significant difference (P<0.05). ± SEM. FM – Fish Meal, SBPC – Soybean Protein Concentrate meal, SBPC<sup>AA</sup> – SBPC supplemented with methionine and lysine.

## 4. Discussion

Results indicate that substituting FM with up to 40 % SBPC in the Nile tilapia diet, replacing 80 % of fish protein did not show significant differences in the investigated parameters concerning growth or carcass composition when compared to the control diet. This study indicates that a 40 % fishmeal replacement could effectively support fish without compromising growth and feed utilisation efficiency. Elangovan and Shim, (2000) showed similar findings with tin foil barb (*Babodes altus)*. Their results confirmed soybean meal could be included at up to 37 % of the diet, replacing approximately 33 % of the fish protein without significantly affecting the growth rate. However, it should be noted that tilapia is an omnivorous species and should be able to tolerate much higher inclusions of plant proteins as a main component of the diet, as explained by Jauncey, (1998) in this author’s comprehensive treatise on tilapia feeds and feeding.

For Anti-Nutritional Factors (ANFs) found within soybeans, it is required to expose soybeans to heat treatment processes before utilisation for monogastric animals. For example, this reduces the concentration of protease inhibitors such as trypsin inhibitors (González-Vega *et al*., 2011). However, excessive heating may also reduce protein and amino acid digestibility and bioavailability, thus negatively impacting the fish’s growth performance. The soybean meal concentrate used in the current trial had been effectively heat treated to denature specific ANFs such as trypsin inhibitor proteins. Another limiting factor in soy-based diets is the lower amounts of Essential Amino Acids (EAAs) compared to fishmeal, such as methionine and lysine, which can be the first and second limiting. Methionine is considered the first limiting essential amino acid in some Nile tilapia-fed cereal-based diets, particularly soybean-meal-rich diets. Supplementation of feed-grade lysine has also been largely adopted in practical and experimental diets for Nile tilapia. Lysine’s primary metabolic role in protein synthesis is also the basis for it to be the reference Amino Acid (AA) in computing ideal AA ratios. Interestingly, an early study demonstrated that the effectiveness of using intact lysine from high-lysine corn protein concentrate was not significantly different from that of crystalline lysine in Nile tilapia (Nguyen and Davis, 2016). He *et al*., (2013) established the methionine and lysine requirements for maintenance and efficiency of utilisation for growth of two sizes of tilapia (*Oreochromis niloticus*). Prabu *et al*., (2021) also worked on tilapia to examine the supplementation of key crystalline amino acids, namely methionine, lysine, tryptophan, and arginine. These researchers used a high soybean meal formulation (analogous to the current study) to assess the respective essential amino acid requirements by their individual deletion from the reference diet. Interestingly, tilapia did not require added tryptophan in their study. These latter authors obtained maximum performance in balanced diets with essential amino acid supplementation (methionine, lysine, and arginine) compared to an un-supplemented group.

In the present study, the supplementation with the two EEAs, methionine, and lysine, resulted in improved growth rates for the higher soybean meal inclusion level. The 80SBPC diet, containing no additional amino acids and the highest soybean protein concentrate inclusion, resulted in the poorest FCR and SGR. On the other hand, the 80SBPC^AA^ diet, supplemented with the EEAs, showed better FCR and SGR in relation to 80SBPC fed tilapia. Furthermore, Webster *et al*., (1995) suggested that a diet that consisted purely of plant protein (e.g. soybean meal) could replace a fishmeal diet without adverse effects, providing methionine was supplemented in the diet. Hu *et al*., (2008) conducted a similar study on gibel carp *(Carassius gibelio)* which investigated diets with and without EAAs lysine and methionine supplementation. The results illustrated the best SGR and final body weight when fish were fed a diet containing lysine and methionine supplementation. This can be seen in the 80SBPC^AA^ diet compared to the 80SBPC non-supplemented group. Crystalline Amino Acids (CAAs) can successfully be included at 60 g kg^−1^ in the Nile <u>tilapia</u> diet as stated by Da Cruz *et al*., (2021). CAAs can stimulate protein synthesis coupled to lower fat deposition in Nile tilapia. The mRNA levels of *mTOR* and *Raptor* genes are up-regulated by proper CAA supplementation. Crystalline amino acids could be used as a strategy to improve amino acid balance beyond elaborating eco-friendly diets. This may explain the superior protein retention found in the present study in terms of ANPU and carcass retention of protein.

Recently, Furuya, Cruz and Gatlin (2023) reviewed the essential amino acid requirements for tilapia, focusing on precision protein nutrition and mentioning the role of supplementary EAAs in diets for optimum performance. The carcass ash content was found to be reduced progressively between diets with increased soybean supplementation and significant only at 80SBPC. This may be due to the declining mineral content associated with increasing soybean meal inclusion. Although we have adjusted for the phosphorous and calcium by adding supplementary inorganic salts as a mineral premix, the phytic acid within soybean chelates these minerals, especially phosphorous within the diet. This leads to a lower availability of the minerals to the fish and, consequently, the reduced concentrations of ash noted in the carcass.

It should be noted that, in general, tilapia all responded well to each diet treatment, and growth rates were comparable to previous work on tilapia by Bell and Davies (2024) and with comparable feed utilisation parameter data obtained from this other study by the same authors. We show that up to 40 % SBPC can replace FM without any significant detrimental effects on growth or carcass composition. Comparison between the data obtained for the 40SBPC diet and the 80SBPC diet, respectively, indicates that there is a slight difference in carcass composition, notably protein and hence growth. It, therefore, indicates that the component nutrients, for example, crude protein and lipids, were not fully bioavailable to the action of digestive enzymes in the tilapia. The previous findings of Lin and Luo (2011) working with juvenile tilapia supports such a view. These authors stated that the higher the dietary inclusion of soybean meal, the lower the activity of proteases in the intestine and hepatopancreas and these were constraining to digestive efficiency.

In summary, our investigation demonstrates soybean protein concentrate can be used as a contributory protein source in diets for Nile tilapia up to a 40 % dietary inclusion threshold. Beyond this level, the soy diet appears deficient in available energy and amino acids, notably lysine and methionine. This absence is detrimental to fish growth and carcass composition. Supplementation with coated crystalline amino acids can, to some degree, correct for the imbalance in soybean meal inclusion at higher levels. The optimal supplementation of crystalline amino acids in feeds for fish provides an opportunity to reduce formulation costs in a quite volatile commodity market of protein ingredients and supply of fish meals. Further research is warranted on determining the potential of using supplementary amino acids in alternative plant-based ingredients in diets for tilapia and other warmwater species. This should be undertaken for different-sized fish during the various production stages and towards harvest within finishing diets, where protein levels are more limiting for essential amino acids.

## Acknowledgements

The authors are grateful to Tom Corry and Wayne Jackson for the practical aspects of this project and technical support assistance for the feeding trials.

## Ethical approval

The animal husbandry techniques involved in the study were undertaken under the supervision of Professor Simon Davies in accordance with the Animal (Scientific Procedures) Act of 1986, License #Pil: 30/00080

## Conflict of Interest

The authors declare that there is no conflict of interest relating to this research study.

## Notes

### Competing Interest Statement

The authors have declared no competing interest.

